# Missense mutations in Myc Box I influence MYC cellular localization, mRNA partitioning and turnover to promote leukemogenesis

**DOI:** 10.1101/2023.10.22.563493

**Authors:** Nancy BJ Arthur, Keegan A Christensen, Kathleen Mannino, Marianna B. Ruzinova, Ashutosh Kumar, Agata Gruszczynska, Ryan B. Day, Petra Erdmann-Gilmore, Yiling Mi, Robert Sprung, Conner R. York, R Reid Townsend, David H. Spencer, Stephen M. Sykes, Francesca Ferraro

## Abstract

Somatic missense mutations in the phosphodegron domain of the *MYC* gene (<u>M</u>YC Box I) are detected in the dominant clones of a subset of acute myeloid leukemia (AML) patients, but the mechanisms by which they contribute to AML are unknown. To unveil unique proprieties of MBI MYC mutant proteins, we systematically compared the cellular and molecular consequences of expressing similar oncogenic levels of wild type and MBI mutant MYC. We found that MBI MYC mutants can accelerate leukemia by driving unique transcriptional signatures in highly selected, myeloid progenitor subpopulations. Although these mutations increase MYC stability, they overall dampen MYC chromatin localization and lead to a cytoplasmic accumulation of the mutant proteins. This phenotype is coupled with increased translation of RNA binding proteins and nuclear export machinery, which results in altered RNA partitioning and accelerated decay of select transcripts encoding proapoptotic and proinflammatory genes. Heterozygous knockin mice harboring the germline MBI mutation *Myc p.T73N* exhibit cytoplasmic MYC localization, myeloid progenitors’ expansion with similar transcriptional signatures to the overexpression model, and eventually develop hematological malignancies. This study uncovers that MBI *MYC* mutations alter MYC localization and disrupt mRNA subcellular distribution and turnover of select transcripts to accelerate tumor initiation and growth.

## INTRODUCTION

MYC (also known as c-MYC) is a multifunctional transcription factor thought to transcriptionally regulate 15% of all known coding genes (Dang et al. 2006; Meyer and Penn 2008). As such, MYC has been implicated in a variety of cellular processes in both malignant and non-malignant conditions, including cell cycle, apoptosis, energy production, anabolic metabolism, cell growth, DNA replication and ribosome biogenesis (Dang 1999; Dang et al. 2006; Posternak and Cole 2016; Dhanasekaran et al. 2022). While *MYC* RNA and protein have short half-lives and are tightly regulated under normal circumstances, MYC expression is frequently dysregulated in a variety of cancers (Dang 2012; Jung et al. 2017).

The MYC protein consists of an amino-terminal transactivation domain, a central intrinsically disordered region, and a carboxy-terminal DNA-binding domain (Murre et al. 1989; Gregory and Hann 2000; Conacci-Sorrell et al. 2014a). MYC contains six conserved motifs called MYC boxes (MB 0-I-II-IIIa-IIIb and IV) (Atchley and Fitch 1995). MB0, MBI, and MBII are located within the transactivation domain, which is required for the transcription initiation, elongation, and the cell transforming activities of MYC (Kato et al. 1990). Within MBI, the phosphorylation of Threonine 73 (T73) by the growth-regulated kinase 3 beta (GSK3β) leads to MYC ubiquitination and degradation via FBW7 (Welcker et al. 2004), while the phosphorylation of Serine 77 (S77) by MAP-kinases leads to MYC stabilization and activation (Sears et al. 2000). Mutations in the *MYC* MBI domain have been described in cancers (Smith-Sorensen et al. 1996) and their oncogenic activities have been attributed to their ability to increase MYC protein levels, and consequently MYC activity, within expressing cells by interfering with the ubiquitination process (Salghetti et al. 1999). Although the Thr-73 and Ser-77 residues in MYC Box I influence protein half-life through regulation of ubiquitin-mediated MYC degradation, data from studies using reporter systems indicated that replacement of these phosphorylation sites with different amino acid residues can cause differential effects on the transactivation capabilities of MYC (Gupta et al. 1993), underscoring the complex nature of MYC’s role in transcriptional regulation. To this end, there is a growing appreciation that the function of MYC extends beyond its role as transcription factor (Das et al. 2023). Therefore, continued studies are necessary to gain a more comprehensive understanding of MYC’s activities during malignant transformation.

In AML, *MYC* is commonly upregulated as a downstream consequence of recurrent AML-associated mutations and fusion genes, such as *FLT3*-ITD, *RUNX1-RUNX1T1*, *PML-RARA*, and others (Salvatori et al. 2011), or due to amplification of all or part of chromosome 8, where *MYC* is located (Jones et al. 2010; A et al. 2018). While missense *MYC* mutations have been detected in only ∼2% of *de novo* AML cases (Papaemmanuil et al. 2016; Tyner et al. 2018), we became interested in studying these mutations after observing their enrichment in normal karyotype AML patients who achieved long-lasting leukemia remissions after treatment with cytarabine-based chemotherapy regimens (Ferraro et al. 2021). To date, the mechanisms by which MYC contributes to myeloid leukemogenesis remains understudied. For instance, it is not known whether *MYC* missense mutations alter hematopoietic stem and progenitor cell (HSPC) function and/or promote AML. Moreover, the consequences of overexpression or mutation of MYC on chromatin binding and transcriptional output in myeloid hematopoietic progenitor cells remains uncharacterized.

Here, we employed doxycycline responsive vectors to study the cellular and molecular consequences of regulated expression levels of MYC^WT^ or MBI MYC^MUT^ proteins in murine hematopoietic stem and progenitor cells. Our results show that when expressed at similar levels, MBI MYC^MUT^ induce a myeloid leukemia *in vivo* with reduced latency compared to MYC^WT^. Like oncogenic MYC^WT^ overexpression, MBI MYC^MUT^ upregulate the expression of self-renewal signatures in hematopoietic progenitor cells. However, MBI MYC^MUT^ display unique proprieties characterized by lower apoptotic rates, and a relative downregulation of pro-apoptotic and pro-inflammatory pathways in distinct and highly selected populations of lineage-restricted granulocyte–macrophage progenitors (GMPs). Notably, many of the transcriptional changes that are unique to MBI MYC^MUT^ expressing GMPs mirror the transcriptomic profiles of AML cells from patients carrying *MYC* mutations. Chromatin immunoprecipitation sequencing (ChIP-seq) analyses of MBI MYC^MUT^ expressing myeloid progenitor cells demonstrate an overall lower DNA binding signal compared to similar oncogenic levels of MYC^WT^ expression which may be due to aberrant cytoplasmic accumulation of the mutant proteins. Results from nascent proteomic analyses in myeloid progenitor cells, uncovered that the induction of MBI MYC^MUT^ (but not of MYC^WT^) proteins promotes the preferential translation of regulators of cell cycle progression, RNA nuclear export, RNA metabolism, which results in subcellular redistribution of mRNA transcripts and accelerated decay of selected mRNA in MBI MYC^MUT^ expressing cells. To validate these findings under physiological levels of expression of MBI MYC^MUT^, we generated mice carrying the germline *Myc* mutation *p.T73N*, a common MBI mutation detected in both myeloid and lymphoid tumors, in the endogenous *Myc* locus. Similar to the overexpression model, *MycT73N/+* mice have cytoplasmic-predominant localization of the MYC protein compared to wildtype littermates, develop bone marrow expansion of GMP-like progenitor cells with similar transcriptional phenotypes and ultimately progress to develop exclusively hematopoietic tumors.

These findings highlight that increases in MYC protein stability and levels—a hallmark of MYC-driven tumors— can have complex and non-linear effects on MYC’s transcriptional activities and suggest that MBI *MYC* mutations can act in a non-canonical fashion to influence RNA partitioning, stability, and protein translation to promote leukemogenesis. Furthermore, the knockin mouse phenotype suggests that MBI MYC mutations are driver mutational events in hematopoietic tumors.

## RESULTS

### MYC BOX I (MBI) *MYC* mutations decrease the apoptotic rate of murine HSPCs leading to an accelerated myeloid leukemia development compared to similar levels of wildtype MYC overexpression

Somatic *MYC* mutations were first recognized as pathogenic events in AML in the context of cryptic *NUP98-NSD1* translocations (Lavallee et al. 2016). In a larger sequencing study, missense *MYC* mutations were detected in 28 out of 1540 (1.8%) sequenced *de novo* normal karyotype AML cases (Papaemmanuil et al. 2016). The average variant allele frequency (VAF) of *MYC* mutations of these cases was 36%, suggesting that most AML-associated *MYC* mutations occur in the dominant malignant clone at presentation.

We recently observed an unexpectedly high incidence of *MYC* mutations in a subset of normal karyotype AML patients who achieved durable remissions after treatment with standard-of-care chemotherapy regimens containing high-dose cytarabine (6/28 cases, 21% vs. 28/1540, 1.8%, Fisher’s exact p<0.0001) (Ferraro et al. 2021). These missense *MYC* mutations also occurred at high VAF (average VAF 38.9%, median VAF 42.8%, range 19.4% - 48.2%, **Suppl. File 1**), and were localized in a discrete, evolutionary conserved MBI region (corresponding to AAs 60-78) (**Suppl. Figure 1A**). Retrospective analysis of AML samples in the COSMIC database corroborated the observation that 80% of all AML-associated *MYC* mutations alter amino acid residues 73, 74 and 75 within MBI (**Suppl. Figure 1B, Suppl. File 1**), which is a pattern that is unique to hematologic malignancies and suggest that this region of MYC may play a critical role in regulating hematopoietic cells.

Because MBI *MYC* mutations are expected to promote oncogenesis by increasing MYC cellular levels, to differentiate dose effects (related to overall MYC levels), from gain-of-function effects (related to the unique properties of each mutant protein), we employed a recombinant retroviral delivery system to model the increase of both wildtype *Myc* (*Myc^WT^*) and mutant *Myc (Myc^MUT^)* in lineage depleted, primary murine hematopoietic stem and progenitor cells (HSPCs). We modeled various point mutations in the two most frequently mutated MBI amino acid residues (p.P74 and p.T73) and in two non-MBI amino acids (p.A59 and p.E137), as detected in our previous study (Ferraro et al. 2021). Initial western blot analysis revealed that HSPCs transduced with either *Myc^WT^* or AML-associated *Myc^MUT^* led to a comparable increase of MYC steady state protein levels compared to empty vector (EV) controls (2 to 5 fold average increase over EV controls, **Suppl. Figure 1C**).

Because the MBI region is known to regulate MYC protein degradation (Conacci-Sorrell et al. 2014a), we performed a cycloheximide pulse-chase experiment to determine MYC^MUT^ vs. MYC^WT^ proteins half-lives. From this analysis, we found that MBI MYC^MUT^ proteins, but not *non-*MBI MYC^MUT^ proteins, displayed significantly longer half-lives compared to MYC^WT^ (**Figure 1A**). We next defined how each *Myc* mutation affected the growth of HSPCs *in vitro*. At 48 hours post-transduction, fluorescent activated cell sorting (FACS) purified GFP+ cells from each condition were assessed for cell expansion every 6 hours for 4 days. This analysis showed that HSPCs transduced with MBI *Myc^MUT^* displayed a statistically significant (P<0.001 at day 4) growth advantage over HSPCs transduced with *EV*, non-MBI *Myc^MUT^*or *Myc^WT^* (**Figure 1B**).

**Figure 1.**
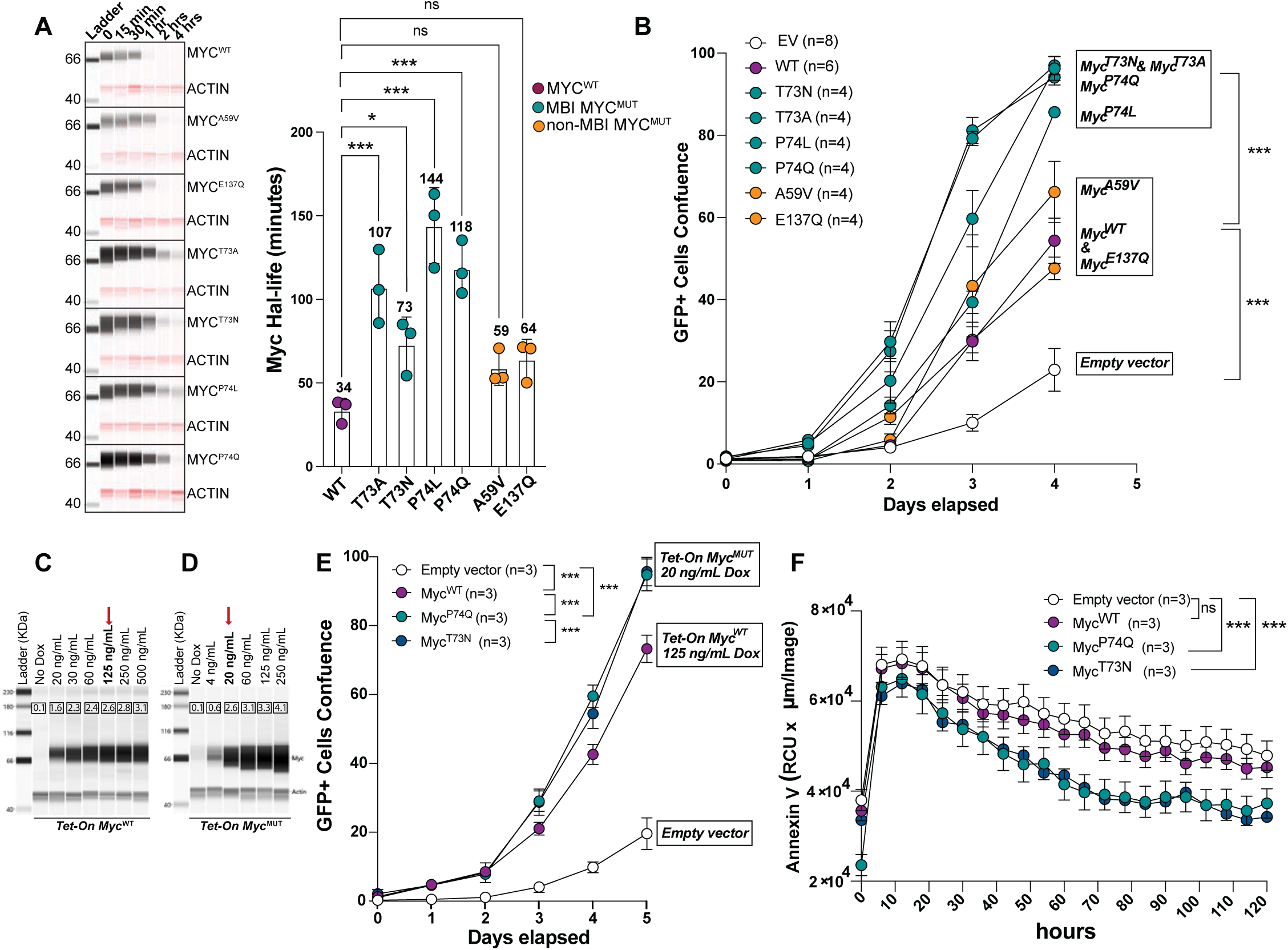
AML-associated *MYC* mutations *in vitro* phenotypes. (**A**) 1×10^6^ GFP+, lineage depleted (Lin^low^) bone marrow cells isolated from C57Bl6 mice, and transduced with wild-type and mutated *Myc* vectors were lysed after treatment with Cycloheximide after 0 (controls), 15 min, 30 min, 1 hr and 2 hrs and subjected to western blot analysis using the JESS automated system. <u>Left panel</u> Representative lane-view images showing MYC and ACTIN bands in lysates from cells expressing WT and various MUT MYC proteins. <u>Right panel</u>: Histogram plot showing the average and standard deviation of MYC half-life for WT and MUT proteins. N=3 for all constructs. Mean half-life is indicated above each bar (*p<0.05, ***p<0.001) (**B)** Expansion (GFP+ cells confluence) of GFP^+^Lin^low^ cells expressing wildtype MYC vs. selected MYC mutants vs. Empty Vector (EV) using the IncuCyte® S3 Live-Cell Analysis System. The number of biological replicates performed for each construct is shown in the legend (***p<0.001, ****p<0.0001 at day 4). **C**) Western Blots and relative quantification (in rectangles within each lane) of MYC protein in mHSPC transduced with Tet-On *Myc^WT^*, or (**D**) Tet-On *Myc^P74Q^*exposed to different doxycycline concentrations. Red arrows indicate doxycycline concentrations with similar MYC protein levels at the time of analysis. (**E**) 5-days expansion and (**F**) apoptosis (as measured by Annexin V staining) of Lin^low^GFP^+^ murine HSPC transduced with Tet-On *Myc^WT^*, *Myc^P74Q^*, or *Myc^T73N^* at doxycycline concentrations resulting in equivalent MYC protein levels (p<0.001 at day 5); the Empty Vector growth curve is shown for reference.

To investigate whether the observed accelerated cell growth was a direct consequence of MYC protein levels, we generated a doxycycline inducible (Tet-On) vector system that allows for controlled expression of WT or mutant MYC proteins by varying the doxycycline dose, and simultaneously constitutively express the fluorescent protein Venus for cell selection (see Supplementary Appendix). Using this system, we observed that at equivalent levels of steady state MYC (compare MYC^WT^ at 125 ng/ml with MYC^MUT^ at the 20 ng/ml dose, **Figure 1C-D**), the MBI MYC^MUT^ expressing HSPCs continued to display a significant growth advantage over MYC^WT^ (**Figure 1E**). This phenotype was related to a concurrent decrease in apoptosis, as evidenced by reduced Annexin-V positivity over time (**Figure 1F**). This result is consistent with a previously reported resistance to *Tp53* indued apoptosis in cells overexpressing the T73A mutation (Hemann et al. 2005).

Syngeneic recipient mice transplanted with primary HSPCs transduced with recombinant retroviruses expressing *Myc^WT^* develop a fully penetrant myeloid leukemia characterized by leukocytosis, bone marrow (BM) infiltration and splenomegaly (Luo et al. 2005). To determine whether MYC MBI mutations can also promote AML development, we took advantage of this model and focused on further characterizing the *in vivo* consequences of the two most common AML-associated MBI missense mutations (p.P74Q and p.T73N) in primary and secondary recipient mice (**Figure 2A**). We transplanted 0.5 x 10^6^ lineage depleted (Lin^low^) HSPCs cells transduced with *Myc^WT^*, *Myc^P74Q^* or *Myc^T73N^* vectors, each with similar fractions of GFP^+^ cells (range 8-10%) into lethally irradiated syngeneic recipient mice. Mice transplanted with Lin^low^ HSPCs overexpressing *Myc^WT^*developed fully penetrant AML with a median survival of 58 days. In keeping with the *in vitro* growth kinetics, primary mice transplanted with HSPCs transduced with both MBI *Myc^MUT^* vectors developed a phenotypically similar disease, with a shortened median survival of 41 days (**Figure 2B**). Western blot analysis for MYC protein expression in extracts from bone marrow cells derived from leukemic primary recipients showed no significant difference in levels of expression of *MYC^WT^* and *MYC^MUT^*proteins (**Figure 2C**).

**Figure 2.**
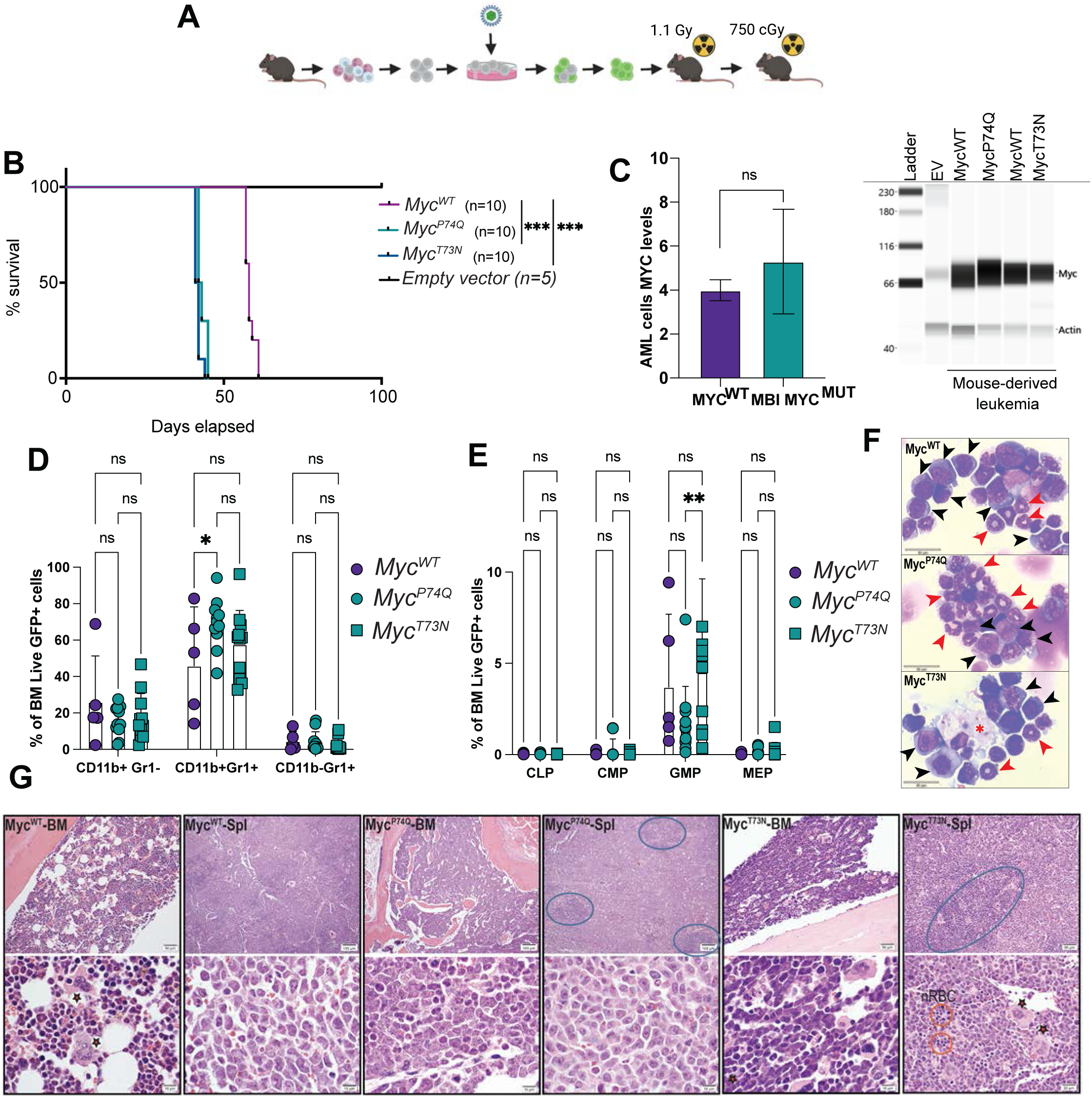
AML-associated MYC mutations in vivo phenotypes. (**A**) Experimental schema for the in vivo experiments: Linlow cells were transduced with either wildtype or mutants Tet-on Myc vectors, then transplanted into lethally irradiated recipients, that were maintained on doxycycline chow. Leukemia cells from each primary mouse that developed disease were then transplanted into secondary, sub-lethally irradiated recipients. (**B**) Kaplan-Meier curves showing overall survival of primary recipients transplanted with HSPCs transduced with MycWT, MycP74Q, MycT73N or Empty Vectors. The number of mice transplanted in each arm is indicated in the legend (***p<0.001, ns=not significant). (**C**) Western Blot and relative quantification of MYC levels in BM cells isolated from mouse derived AMLs transduced with MycWT, MycP74Q, MycT73N and EV control BM cells (ns=not significant). (**D and E**) Histogram plots of multiparameter flow cytometry data of bone marrow derived cells from sick mice at the time of takedown. The majority of GFP+ cells expressed cell surface markers of either (**D**) maturing myeloid cells or (**E**) granulocyte/macrophage progenitor (GMP: Linlow, Sca-1-, c-kit+, CD34+ and CD16/32+) (columns are mean ± standard deviation, *p<0.05, **p<0.01, ns=not significant). (**F**) Cytospin images showing the myeloid morphology of the tumors derived from primary transplant recipients. Note the presence of immature-looking, large cells with round nuclei, nucleoli, and cytoplasmic granules (black arrowhead) and the presence of smaller, neutrophil-like cells with C-shaped or donut-like nuclei (red arrowhead). The star indicates a macrophage. Scalebar: 20 microns (µm). (**G**) Hematoxylin-Eosin of bone marrow and relative spleen sections from sick mice transplanted with Linlow cells expressing WT, P74Q or T73N MYC proteins. The bone marrow in mice expressing MYC wildtype appears relatively preserved compared to the mice expressing MYC mutants. The spleens in all three cases are hypercellular and involved by sheath of large mononuclear cells that resemble blasts on a background of myeloid elements (better appreciable in the bottom panels, which represent higher magnification details of each top panel). Note that the parenchyma of the spleens is mostly effaced by atypical cells with vesicular chromatin and prominent nucleoli. Increased, clustered, atypical megakaryocytes are also appreciable, which appear to be particularly prominent in the MycT73N leukemias (highlighted by red stars). Nucleated red blood cells are also visible (red circles). The areas delimited by blue circles represent occasional residual normal splenic parenchyma. Scalebar: 100, 50 or 10 microns (µm), as indicated in each panel.

Secondary recipient mice transplanted with 1.0×10^6^ bone marrow-derived primary leukemia cells expressing either MYC^WT^ or MBI MYC^MUT^ also developed a fully penetrant leukemia, with similar latency (median survival of 40 days and 45 days respectively, **Suppl. Figure 2A**).

Multiparameter flow cytometric and morphologic analyses of BM cells isolated from primary recipient mice showed an aberrant expansion of GFP^+^ cells expressing myeloid markers with all three vectors (*Myc^WT^*, *Myc^P74Q^*and *Myc^T73N^*, **Figures 2D-F**). Histological analysis of hematopoietic cells from the tibias and spleens (**Figure 2G**) of moribund mice showed the presence of sheets of large, atypical cells variably infiltrating the bones and/or the spleens of sick mice. In the spleens, these infiltrates were evident on background of extramedullary hematopoiesis and atypical megakaryocytes, supporting the myeloid origin of these tumors. There were no significant differences in total WBC counts or spleen weights among *Myc^WT^*, *Myc^P74Q^* and *Myc^T73N^* leukemic mice (**Suppl. Figures 2C-D**). Flow cytometric analysis of peripheral blood (PB), BM, and spleen cells showed a similar distribution of GFP^+^ cells in each compartment across all three genotypes (**Suppl. Figures 2E-F**). Collectively, these data suggest that compared to similar levels of oncogenic MYC^WT^ expression, MBI MYC^MUT^ proteins accelerate the initiation of a frank myeloid leukemia in mice.

### MBI *MYC* mutations drive unique transcriptional signatures in subpopulations of pre-leukemic Granulocyte-Macrophage Progenitor cells

To gain insights in the mechansims related to the accelerated leukemia onset observed in MBI MYC mutant expressing HSPCs, we next used our retroviral in vitro model system to examine the transcriptional programs of HSPCs immediately following expression either *EV*, *Myc^WT^*, *Myc^P74Q^* or *Myc^T73N^*. To gain cellular resolution and to account for changes in cellular populations that may have occurred between transduction and sequencing, we performed single-cell RNA sequencing (scRNA-seq) of lineage depleted (Linl^ow^) HSPCs 48 hours after transduction.

Figure 3A shows a Uniform Manifold Approximation (UMAP) of single cell normalized expression data from HSPCs expressing *Control-EV*, *Myc^WT^,* and each of the two MBI *Myc^MUT^*. Interestingly, analysis of transcriptionally-defined cell populations identified a striking shift in the Granulocyte-Macrophage Progenitor (GMP) compartment across the samples, which was significantly expanded in cells expressing MYC^WT^ and MBI MYC^MUT^ (Figure 3B). Given that we also observed expansion of GMPs in the mouse-derived leukemias (shown in **Figure 2D**), we focused our analyses on these cells (GFP+GMPs). We first examined the shared transcriptional phenotypes derived from expressing oncogenic levels of MYC^WT^ and MBI MYC^MUT^. To do this, we performed a differential expression analysis (ANOVA: log FC>±2, FDR<0.05) comparing *EV* vs *Myc* overexpressing cells (*Myc^W^*^T^ and *MBI Myc^MUT^*) GFP+GMPs. With these criteria, we detected 858 new DEGs (ANOVA, log FC>±2, FDR<0.05, **Suppl. File 2**). Among these, 472 genes were upregulated (cluster 1) and 386 were downregulated (cluster 2) in *Myc* overexpressing cells vs. *EV* controls (**Figure 3C**). To further explore these pathways, we performed a supervised functional enrichment analyses of DEGs using *Enrichr* (Chen et al. 2013; Kuleshov et al. 2016; Xie et al. 2021), and then visualized the results with the Enrichment Analysis visualization Tool *Appyters (Clarke et al. 2021)* (**Figure 3D**). Each of the points in the Bokeh plot represent an annotated gene set from the KEGG 2019 mouse pathway. In this plot, KEGG terms with similar gene sets are clustered together, and are of the same color. Large dots represent significantly enriched gene sets in the input gene list, with the top 10 significantly enriched gene sets represented by dots outlined in black (top 10 pathways). This analysis revealed a clear similarity of the genes sets influenced by *Myc* overexpression, indicated by the strong and distinct clustering of each set of upregulated and downregulated genes (**Figure 3D**). Specifically, we found that genes upregulated in *Myc* overexpressing cells were enriched for pathways associated with RNA biogenesis, nucleocytoplasmic transport, RNA stability, nucleotide metabolism, and amino acid metabolism. Conversely, hematopoietic cell lineage differentiation and immune response pathways were significantly downregulated by *Myc* overexpression (**Suppl. File 2**).

**Figure 3.**
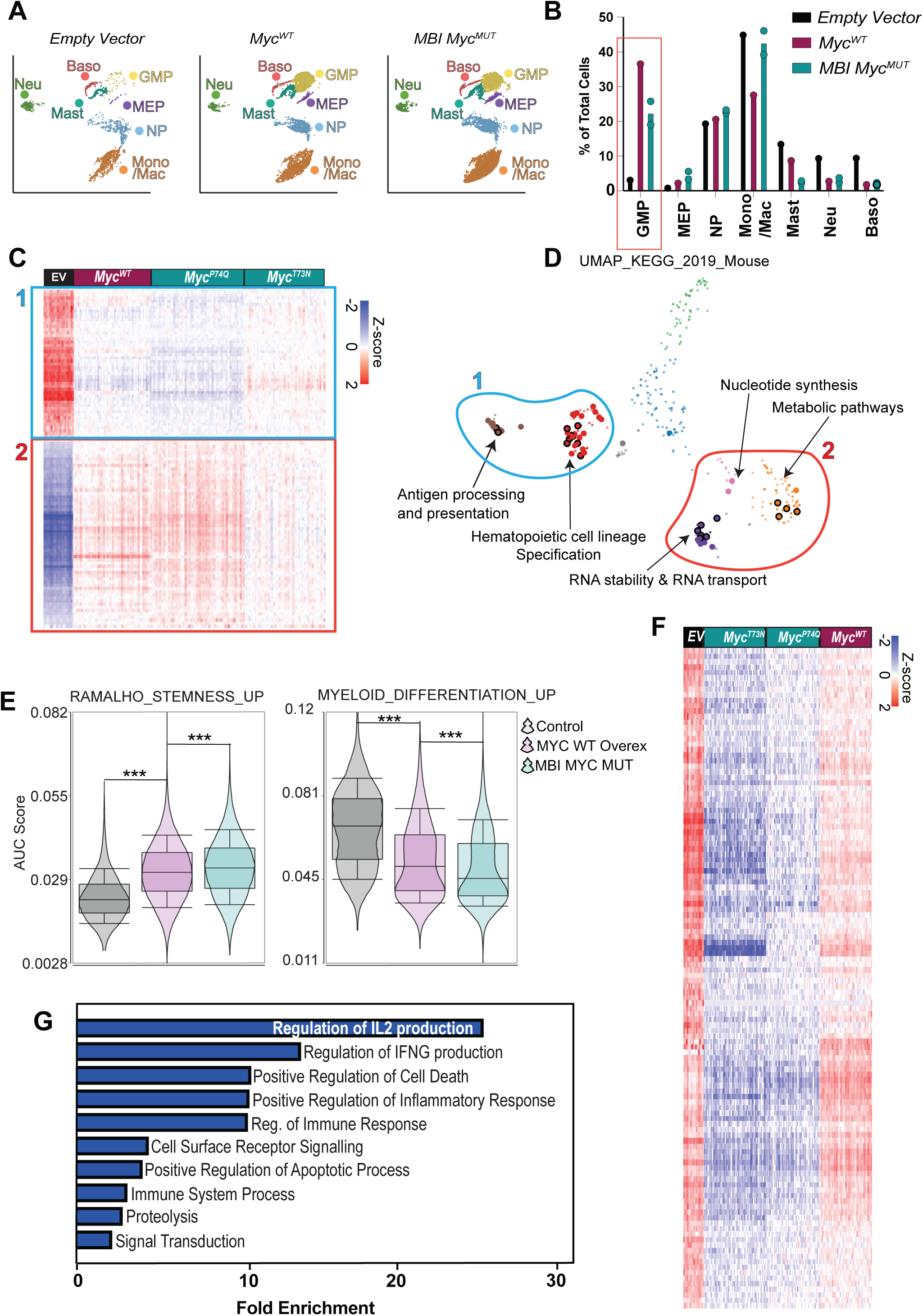
Single cell RNA sequencing studies of preleukemic HSPCs expressing wildtype or mutant MYC proteins. (**A**) UMAP projection of 34985 Lin^low^ GFP+ cells, transduced with *EV*, *Myc^WT^, Myc^P74Q^* and *Myc^T73N^* (combined in *MBI Myc-MUT*) split by genotype and colored by cell lineage (Neu= neutrophils, Baso=basophils, GMP=granulocyte macrophage progenitor, MEP= megakaryocyte erythroid progenitor, NP= neutrophil precursor, Mast=mast cells, Mono/Mac=monocyte macrophages progenitors). (**B**) Bar graph representing the cell type distribution as percentage of total cells in each genotype. (**C**) Heatmap representing the DEGs in GFP+ GMP cells of EV vs. Myc overexpressing cells (wildtype + mutant) The Empty Vector control is plotted passively. (**D**) Scatterplot of all terms in the KEGG_2019_Mouse gene set library. Each point represents a term in the library. Term frequency-inverse document frequency (TF-IDF) values is computed for the gene set corresponding to each term, and UMAP is applied to the resulting values. The terms are plotted based on the first two UMAP dimensions. Generally, terms with more similar gene sets are positioned closer together. Larger and outlined dots represent the top significantly enriched terms. (**E**) Area under the curve (AUC) score for gene lists associated with stemness (left panel) or myeloid differentiation (right panel), in *EV*, *Myc^WT^* and *MBI Myc^MUT^* single cells. ***=FDR<0.001 (**F**) Heatmap representing the DEGs in GFP+ GMP cells of *Myc^WT^* vs. *MBI Myc^MUT^* overexpressing samples. (**G**) Enrichment for Gene Ontology terms in the DEGs represented in (**F**). Blue bars are significantly (FDR<0.05) downregulated pathways by MBI *Myc* mutations.

Integrated expression analysis of publicly available gene lists (Ramalho-Santos et al. 2002; Terskikh et al. 2003), showed that genes implicated in hematopoietic stem cell self-renewal were significantly enriched in HSPCs expressing MBI MYC^MUT^ vs both MYC^WT^ and EV, whereas genes involved in myeloid differentiation and lineage commitments were reciprocally downregulated (**Figure 3E**). This was associated with an increase in clonogenicity of pre-leukemic myeloid progenitors in MBI MYC^MUT^ versus MYC^WT^, as evidenced by the higher colony formation with subsequent passages in cytokine-enriched methylcellulose stem cell media (**Suppl. Figure 3A**).

Next, we focused on transcriptional programs unique to MBI MYC^MUT^. By employing ANOVA, we identified 247 differentially expressed genes when comparing *Myc^WT^ vs. MBI Myc^MUT^*GFP+ GMPs cells (log FC>±2, FDR<0.05, **Figure 3F** and **Suppl. File 3**). The heatmap in Figure 3F, shows that these genes are highly expressed in control GMP cells (*EV*), while *MBI Myc^MUT^* GMP cells exhibit the lowest expression values relative to both *Myc^WT^* and *EV* GMP cells.

Consistent with the measured reduction in apoptotic rate *in vitro* and the shortened leukemia latency observed in the MBI MYC^MUT^ expressing cells (**Figures 1F** and **2B**), genes selectively downregulated in MBI MYC^MUT^ GMP cells were enriched of proapoptotic genes (GO biological processes, FDR<0.05, **Figure 3F, Suppl. File 4**). Furthermore, the MBI MYC^MUT^ specific cluster was comprised of immune response and inflammatory chemokines transcripts (**Figure 3F, Suppl. File 4**). Amongst these, the downregulation of *Cd34* and Major Histocompatibility Complex genes (*MHC*) resembled that observed in AML patients with *MYC* mutations, reported in our recent study (Ferraro et al. 2021), where AML cells with *MYC* mutations expressed lower levels of *MHC* and *CD34* transcripts relative to AML cells without *MYC* mutations.

To further examine the association between *MYC* mutations and the expression of the MBI MYC^MUT^- unique downregulated gene cluster in patient-derived AML samples, we calculated the Area Under the Curve (AUC, see supplementary appendix) for the expression of the 247 genes in single cell transcriptomes of MYC^WT^ and MYC^MUT^ AML cells at diagnosis (Ferraro et al. 2021). Consistent with the mouse result, we detected a significantly lower overall expression of this genes in *MYC* mutated human AML cells, compared to AML cells without *MYC* mutations (**Suppl. Figure 3B**).

To validate the mouse scRNA-seq results, we performed two independent bulk RNA sequencing experiments of FACS sorted GFP^+^, Lin^low^ HSPCs from each EV, *Myc^WT^*, *Myc^P74Q^* and *Myc*^T73N^, 48-hours post-transduction, the same time point used scRNA-seq analyses. DESeq2 comparison of EV vs. *Myc* overexpressing cells detected 2824 DEGs (1178 upregulated and 1646 downregulated) in *Myc* overexpressing cells vs *EV* transduced controls (log FC>±2, FDR<0.05, **Suppl. Figure 3C, and Suppl. File 5**). KEGG 2019 pathway enrichment analysis of the bulk RNA-seq DEG overlapped with the KEGG analysis results obtained with the scRNA seq (**Suppl. Figures 3D**). Notably, DEG analysis yielded no differentially expressed genes when comparing Myc-WT vs. Myc-P74Q and only 24 genes when comparing Myc-WT vs. Myc-T73N (DESeq2, with an FDR cutoff of <0.05 and no fold change cutoff. n=2 samples/genotype, **Suppl. File 5**) confirming that the MBI MYC^MUT^ specific changes are unique to highly selected subpopulations of GMP+ cells and cannot be resolved using bulk sequencing techniques.

### MBI MYC^MUT^ has lower chromatin occupancy than oncogenic MYC^WT^ and accumulates in the cytoplasm

We next investigated whether the unique differences in gene expression, accelerated growth kinetics and leukemia initiation capabilities of the MBI MYC^MUT^ expressing cells could be related to differences in transcriptional regulation between the wildtype and mutant proteins. Given MYC’s role as a transcription factor, we next explored the chromatin localization patterns of wildtype and mutant MYC proteins by performing two independent ChIP-seq experiments on Lin^low^ GFP+ sorted, pre-leukemic, primary HSPCs at 48-hours post-transduction (the same timepoint used for the transcriptomic analyses), using anti-MYC antibodies. The results revealed a significant overlap in the ChIP-seq signal between the two biological replicates (67.4% of peaks overlap for MYC^WT^, **Suppl. Figure 4A, Suppl. File 6**). As expected, the distribution of ChIP signals indicated that both MYC^WT^ and MBI MYC^MUT^ primarily localize at or near transcriptional start sites (TSSs), as shown in **Figures 4A-B**. Notably, no significant endogenous signal was detectable in cells transduced with *EV* (**Figure 4B**), confirming that the signal detected in this dataset primarily originated from the exogenous, overexpressed MYC protein.

**Figure 4.**
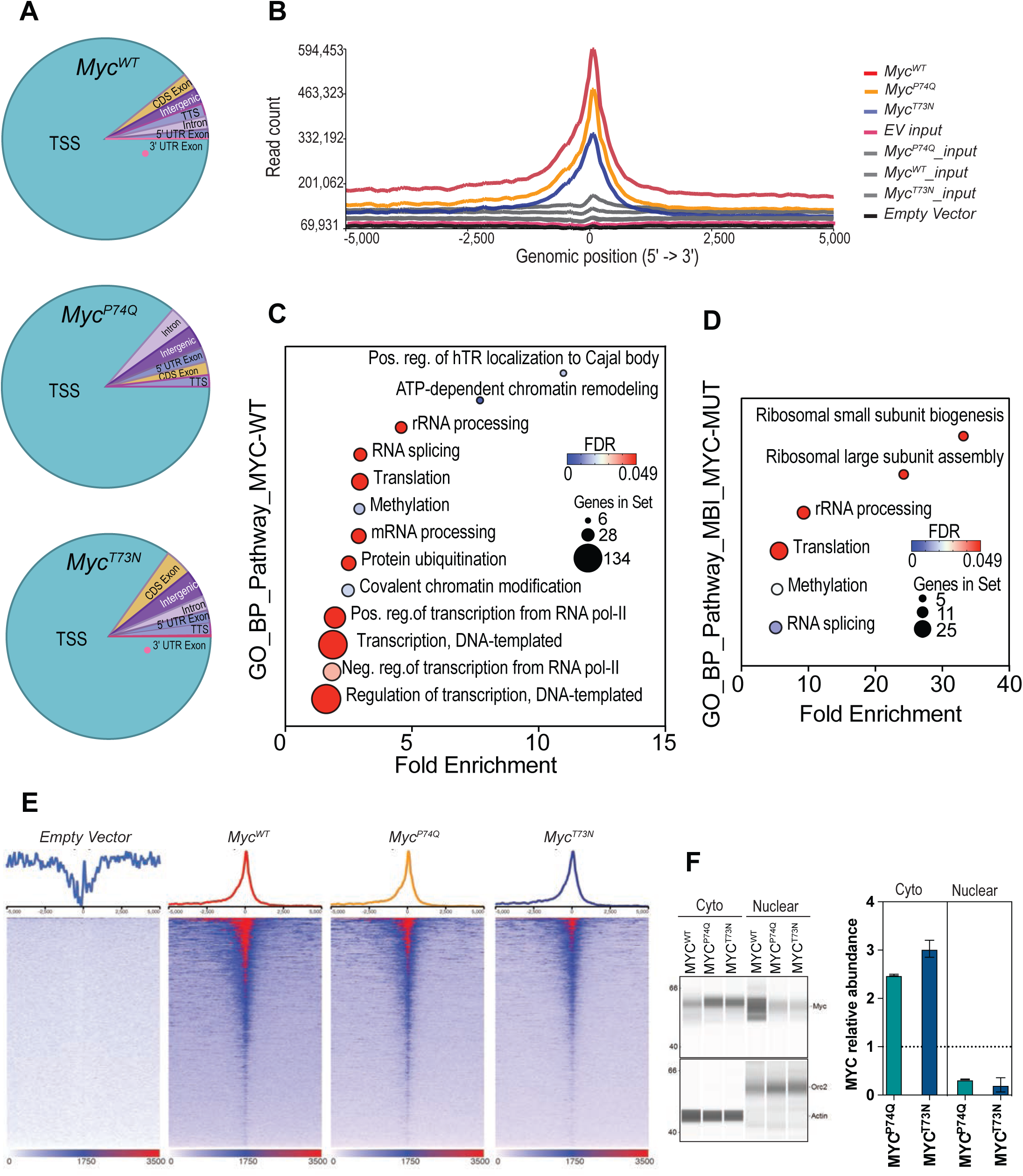
Chromatin binding behaviors of oncogenic MYC^WT^ and MBI MYC^MUT^ expressing cells. (**A**) Pie charts representing the gene section breakdown of ChIP signal derived from MYC^WT^ and MBI MYC^MUT^ proteins (P74Q and T73N). TSS: transcriptional start sites. TTS: transcriptional termination sites. UTR: untranslated region. CDS: coding sequence. (**B**) Distribution of MYC peaks in GFP+ Lin^low^ cells in relation to the nearest transcriptional start site (TSS) in GFP+ Lin^low^ cells transduced with *Myc^WT^, Myc^P74Q^* and *Myc^T73N^* vectors, *EV* (black line) relative to input controls (grey lines). Gene Ontology analysis for MYC-WT ChIP-seq peaks (**C**) and MBI MYC^MUT^ ChIP-seq peaks (**D**). All the pathways represented in the figure are significant by FDR. The size of the circle is proportional to the number of enriched genes within each GO term. (**E**) Heat map of MYC peaks 5Kb upstream and downstream the TSSs for all expressed genes ranked by their signal in GFP+ Lin^low^ cells transduced with *EV*, *Myc^WT^, Myc^P74Q^* and *Myc^T73N^* vectors. The color scale is indicated at the bottom of each heatmap. (**F**) Western blot (left panel) and relative quantification (right panel) of cytoplasmic versus nuclear cellular lysates of GFP+ Lin^low^ cells transduced with *Myc^WT^, Myc^P74Q^* and *Myc^T73N^* vectors. The top bands represent MYC and the lower bands loading controls (ACTIN for cytoplasm and ORC2 for nucleus). The data is represented as fold changes in MYC protein abundance in cells expressing MBI MYC^MUT^ versus MYC^WT^ proteins.

Gene Ontology (GO) analysis of the promoters occupied by MYC^WT^ revealed several pathways consistent with the known functions of MYC, including positive and negative regulation of transcription, chromatin modification, protein ubiquitination, mRNA processing, regulation of translation, and RNA biogenesis (**Figure 4C**). MBI MYC^MUT^ proteins occupied similar transcriptional start sites (including genes involved in mRNA processing and translation) but displayed reduced occupancy at several genes important for myeloid differentiation, regulation of transcription, and ubiquitination (**Figure 4D**). By searching for known DNA-binding motifs in MYC^WT^ and MBI MYC^MUT^ derived peaks, we identified a significant enrichment of the canonical MYC

E-box sequence (CACGTG), indicating that AML-associated *Myc* mutations do not appear to alter the consensus sequence for DNA binding (**Suppl. Figure 4B**). Consistent with this observation, analysis of the ChIP peaks that occurred in gene-coding regions revealed that despite the longer MYC half-life, MBI MYC^MUT^ displayed an overall decrease in DNA occupancy compared to MYC^WT^ overexpression (**Figure 4E, Suppl. Figure 4C**). To ensure that the reduced ChIP-seq signal observed in the MBI MYC^MUT^ cells was not secondary to a reduced affinity of the anti-MYC antibody for the mutant proteins, we used equivalent amounts (5 µg/sample) of the ChIP antibody (rabbit anti-MYC, 9402s, Cell Signaling Technology) to immunoprecipitate MYC^WT^ or MBI MYC^MUT^ from whole cell protein lysates of HSPCs transduced with *Myc^WT^*, *Myc^P74Q^* or *Myc^T73N^* vectors. Immunoprecipitates and inputs from each condition were then subjected to western blot using an independent MYC antibody (mouse anti-MYC, 9E10, sc-40X, Santa Cruz). The amount of MYC protein detected in the precipitates was comparable across all MYC^WT^ and MYC^MUT^ lysates (**Suppl. Figure 4D),** suggesting that the observed differences in promoter localization were not due to a reduced affinity of the anti-MYC antibody for the mutant proteins.

To determine whether the difference in ChIP signal observed between MYC^WT^ and MBI MYC^MUT^ was related to differential cellular localization of the MYC mutant proteins, we performed column-based protein extraction of the cytosolic and nuclear fractions of HSPCs using a commercially available kit (Invent Biotechnologies) 48-hrs after transduction with *Myc^WT^*, *Myc^P74Q^* and *Myc^T73N^* vectors. The results showed that the relative abundance of MBI MYC^MUT^ proteins in the nuclear fraction was significantly lower relative to MYC^WT^. Conversely, MBI MYC^MUT^ proteins were significantly enriched in the cytoplasmic fraction, compared to MYC^WT^, suggesting that these mutations promote MYC accumulation in the cytoplasm, reducing its abundance in the nucleus (**Figure 4F**).

Manual visualization of the 247 promoter regions of genes found to be downregulated in MBI MYC^MUT^ expressing cells using Integrative Genomics Viewer browser (IGV) (Robinson et al. 2011), showed similar signal between wildtype and mutants (see selected examples in **Suppl. Figures 4E-H)**, suggesting that the relative downregulation of these transcripts could not be directly explained by a decrease in promoter regions accessibility or DNA binding of the MBI MYC^MUT^ proteins to these promoter regions.

### MBI MYC^MUT^ influences mRNA partitioning and turnover by enhancing nuclear export

Given that mRNA steady-state levels are a result of an equilibrium between transcription and degradation, to further explore the mechanisms leading to the downregulation of selected transcripts in MBI MYC^MUT^ expressing cells, we next investigated mRNA degradation kinetics in MYC^WT^ vs MBI MYC^MUT^ cells. HSPCs expressing either MYC^WT^ or MBI MYC^MUT^ proteins were treated with 5,6-dichloro-1-beta-D-ribofuranosylbenzimidazole (DRB) at 40 μg/mL, a concentration sufficient to inhibit RNA polymerase II without significantly affecting cells viability (**Suppl. Figure 5A**). Total mRNA was extracted from treated cells at various time points ranging from 0 to 480 minutes. The mRNAs decay rate of cell cycle and immune-regulatory candidates randomly selected amongst the genes downregulated by MBI MYC^MUT^ was determined using qPCR analysis. This analysis showed that the presence of MBI *Myc* mutations shortened the mRNA half-lives of the selected genes when compared to both *Myc^WT^*overexpression and control EV cells (**Figure 5A**).

**Figure 5.**
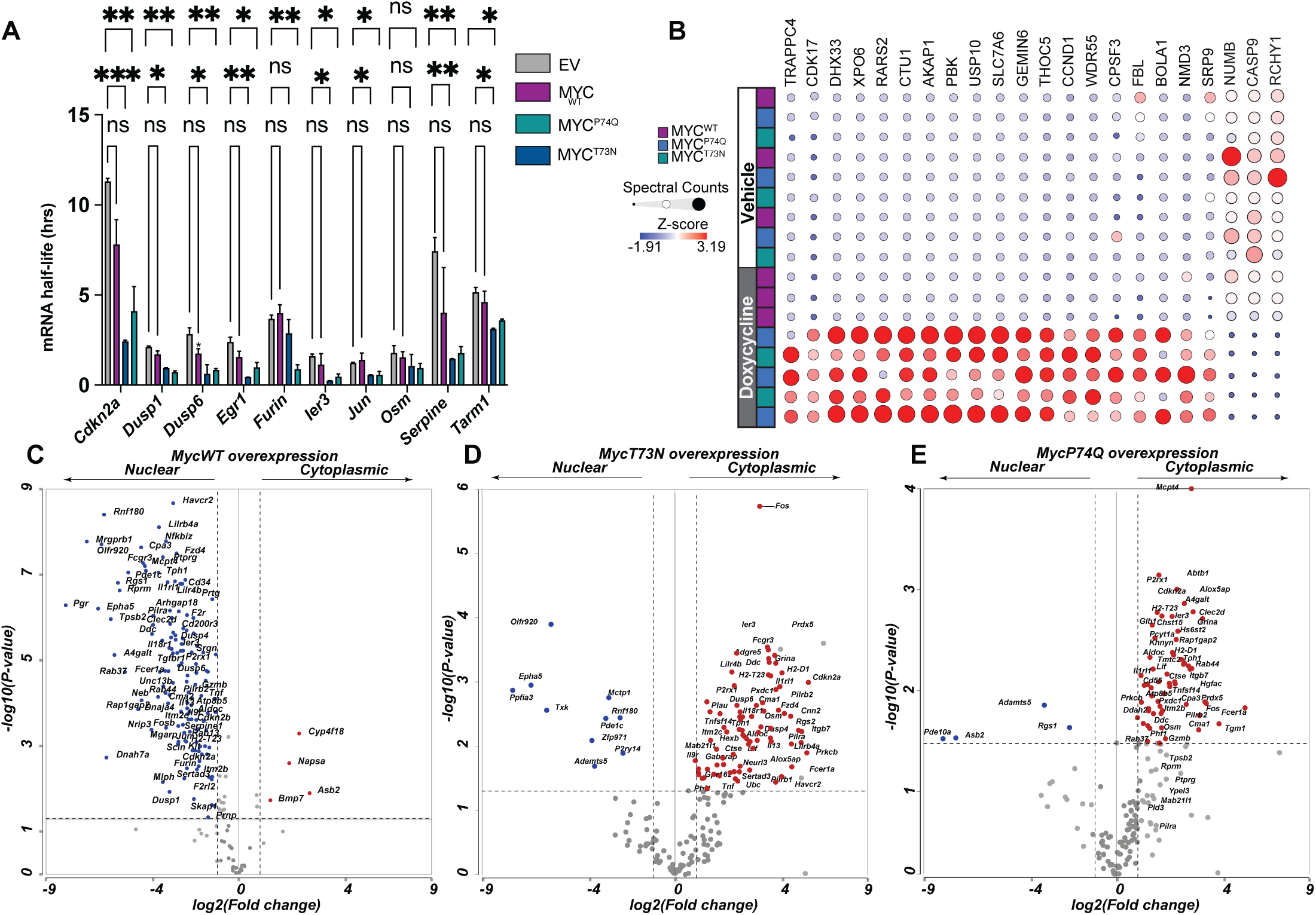
mRNA decay, nascent proteomic analysis, and mRNA nucleocytoplasmic distribution in MYC^WT^ and MBI MYC^MUT^ expressing cells. (**A**) Histogram plots of mRNA half-life as determined by qPCR analysis of selected transcripts in control, MYC^WT^, MYC^P74Q^ and MYC^T73N^ expressing cells. Data are average ± SD, n= 3 biological replicates. (*p<0.05, **p<0.01). (**B**) Unsupervised hierarchical matrix of the average difference (Z-score) in protein expression in MBI MYC^MUT^ vs. MYC^WT^ expressing cells. Z scores are calculated based on normalized spectral counts across 2 independent experiments for inducible (Tet-On) *EV*, *Myc^WT^, Myc^P74Q^*and *Myc^T73N^*. The grey bar indicates doxycycline treatment, the white bar represent vehicle treated cells. (**C-E**) Volcano plots showing nuclear/cytoplasmic ratio changes of mRNA transcripts from Figure 3F (and Suppl. File 3) in HSPCs transduced with (**C**) Myc^WT^, (**D**) *Myc^T73N^ and* (**E**) *Myc^P74Q^*. Differentially distributed transcripts that reach significance are colored in blue when enriched in the nucleus and in red when enriched in the cytoplasm.

To gain insights in the potential mediators of accelerated mRNA decay in MBI MYC^MUT^ expressing cells, we performed nascent proteomic analysis (OPP-ID) and assessed the early changes in translational landscape induced upon expression of wildtype or MBI mutant MYC proteins. Briefly, Lin^low^ HSPCs transduced with *Myc^WT^*or *MBI Myc^MUT^* Tet-On vectors were treated with doxycycline or vehicle for 12 hours and then incubated with O-propargyl-puromycin (OPP) for 30 minutes to pulse-label nascent, actively elongating polypeptides. The labelled peptides were then conjugated onto beads using click chemistry, stringently washed, digested, and analyzed using mass spectrometry (**Suppl. Figure 5B**). From this analysis, we observed that only 22 proteins were differentially translated in HSPCs expressing MBI MYC^MUT^ (19 increased and 3 decreased, DESeq2 of MYC^WT^*Dox vs MBI MYC^MUT^*Dox, adjusted p<0.05, FC>±2, **Figure 5B**). Several of the upregulated proteins upon MBI MYC^MUT^ expression were known mediators of mRNA processing and nuclear export (**Suppl, Figure 5C**). To determine whether these differences affected transcript localization between HSPCs expressing wildtype or mutant MYC, we next employed cellular fractionation followed by RNA extraction and sequencing of nuclear and cytoplasmic mRNA in *Myc^WT^*, *Myc^P74Q^* or *Myc^T73N^*transduced HSPCs. Using DESeq2, we compared the abundance of the MBI MYC^MUT^-downregulated transcripts (adjusted p<0.05 and FC>±2) in the cytoplasmic versus nuclear fractions for each genotype. This analysis revealed that these transcripts were enriched in the nuclear fraction of HSPCs expressing MYC^WT^ (**Figure 5C**) but were depleted in the nuclear fraction (and enriched the cytoplasm) in HSPCs expressing MYC^P74Q^ and MYC^T73N^ (**Figure 5D-E**).

### *MycT73N/+* mice display cytoplasmic-predominant MYC localization and develop hematopoietic tumors

To ensure that our observations were not merely the consequence of supraphysiological expression of MYC proteins, we generated knock-in mice carrying a germline MBI mutation *p.T73N* (**Figure 6A**) in the endogenous *Myc* locus. The selection of the mutation was based on its recurrent occurrence in hematologic malignancies, encompassing both myeloid and lymphoid origin (Bhatia et al. 1993; Hoang et al. 1995). Mice heterozygous for *pT73N* exhibited normal development and fertility. However, attempts to breed heterozygous mice did not yield homozygous pups. Adult, 8-12 weeks old *MycT73N/+* mice displayed normal blood counts, body weight, and distribution of hematopoietic stem and progenitor cells in their bone marrow by flow cytometry (see Suppl. Figure **Suppl. Figure 6B-E**). In line with the anticipated protein stabilization effect, these mice exhibited elevated steady-state levels of MYC protein compared wildtype littermate controls (**Figure 6B**). *MycT73N/+* mice developed hematopoietic tumors, albeit with a long latency (12-15 months) and incomplete penetrance compared to the overexpression model (**Figure 6C**).

**Figure 6.**
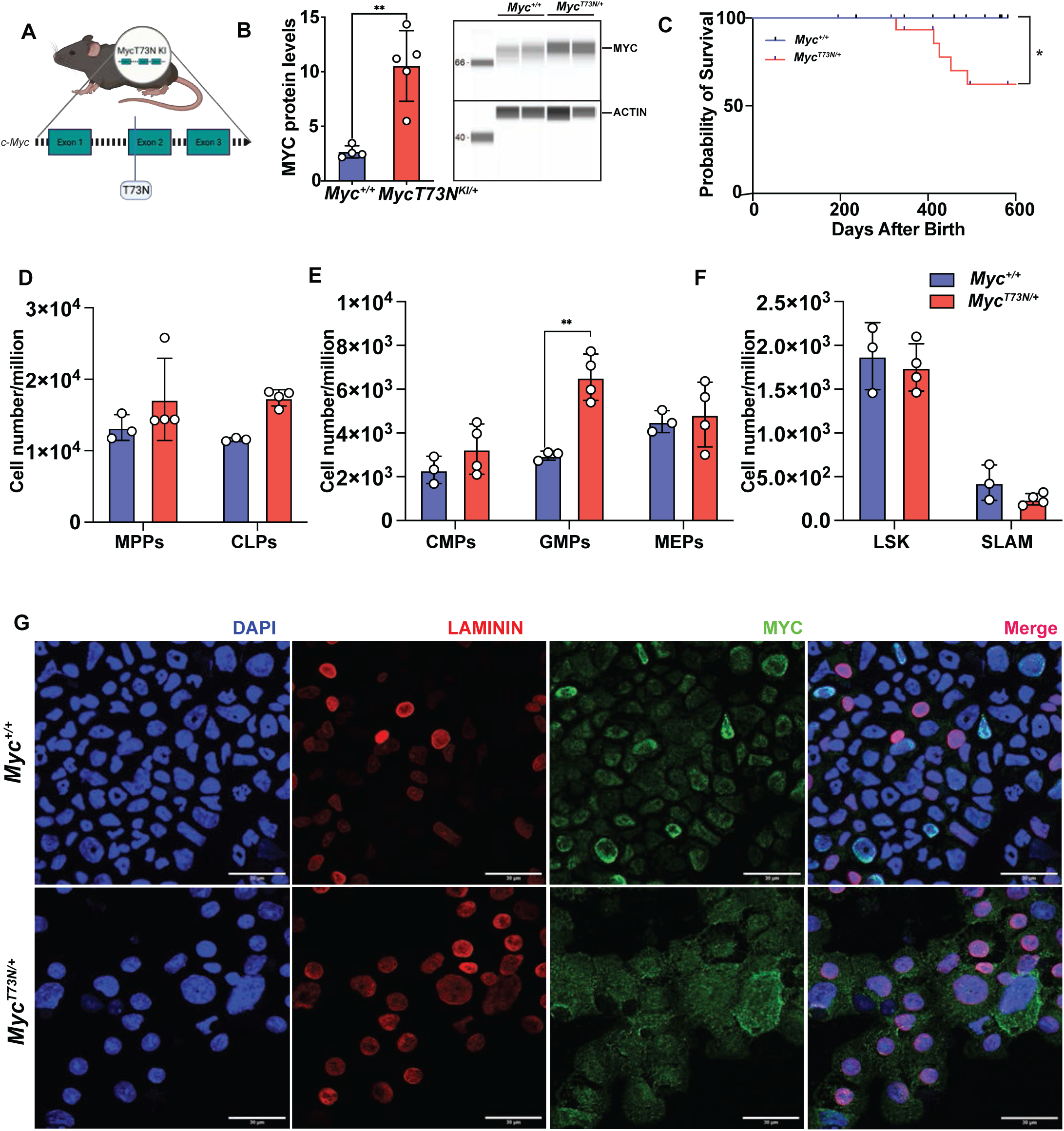
MYC subcellular distribution and survival of MycT73N/+ mice. (**A**) Schematic design of the location of the p.T73N mutation in C57Bl6 mice. (**B**) Histogram plot and representative western blot images showing the steady state levels of MYC protein in HSPCs isolated from MycT73N/+ mice and wildtype littermates (**=p<0.01). (**C**) Kaplan-Meier curves showing the overall survival of MycT73N/+ and wildtype littermates censored at the time of writing (*=p<0.05). (**D-F**) Flow cytometry analysis showing the distribution of hematopoietic progenitor cells in the bone marrow of moribund MycT73N/+ mice at takedown. Age-matched littermate controls were taken down and analyzed side-by-side for comparison (*=p<0,05, all other comparison not significant). (**G**) Confocal images of HSPCs isolated from healthy, 6 months old, MycT73N/+ and littermate controls mice. The blue is DAPI staining (nuclei), the red is laminin A/C staining (nuclear envelope), and the green is MYC staining. Scalebar 30 microns (µm).

Histological examination of femurs from sick mice at takedown revealed hypercellular marrows, with a predominance of myeloid elements (**Suppl. Figure 6F**). Flow cytometry showed absolute expansion of GMP+ progenitor cells (**Figure 6D-F**). Spleens from *MycT73N/+* mice were significantly enlarged compared to control wildtype littermates (*MycT73N/+* average 950 mg ± 513 mg, *Myc^+/+^* average 147 mg ± 24mg, n=5) and contained two abnormal cell populations, which could represent abnormal myeloid and lymphoid infiltrates or, less likely, a biphasic-appearing myeloid population (**Suppl. Figure 6G**). Two out of five animals developed hepatomegaly and lymph nodes enlargement with similar mixed-appearing infiltrates (**Suppl. Figure 6H-I**). Immunofluorescent analysis of bone marrow-derived cells from 8-12-week-old mice showed that, like the overexpression model, MYC localization was predominantly cytoplasmic (**Figure 6G**). These results confirm that endogenous, heterozygous expression of the *p.T73N* missense mutation leads to a cytoplasmic localization of MYC and eventually to the development of bone marrow myeloproliferative disorders, and possibly to mixed myeloid and lymphoid disorders in vivo.

### Pre-leukemic *MycT73N/+* mice display a preferential expansion of a GMP-like population with aberrant transcriptional programs

To explore population specific phenotypes, and to determine the early changes in HSPCs that may be important for the development of leukemia, we performed single cell RNA of bone marrow derived, Lin^low^ HSPCs isolated from pairs of *MycT73N/+* and *wildtype* littermates at 6 months of age. At the time of analyses, these mice appeared healthy, and had no phenotypic evidence of disease. After applying quality metrics filters, we obtained 36085 cells for further analysis. HSPCs cells from pre-leukemic *MycT73N/+* and *wildtype* littermate mice were phenotypically homogeneous and clustered together (**Suppl. Figure 7A**). We next applied graph-based clustering to the data, which parsed the cells in 10 clusters (**Figure 7A**). These clusters were further characterized by differential gene expression analysis with ranking of the top expressed markers in each cluster and deduction of lineage priming using functional enrichment tools (Chen et al. 2007; Chen et al. 2009a; Chen et al. 2009b) (**Suppl. File 7**). Distribution of the different HSPCs populations amongst these clusters revealed that *MycT73N/+* mice had a relative expansion of cluster 1, a GMP-like progenitor population (*Mpo*+, *Elane*+) with a reciprocal decrease of cluster 10, a granulocytopoietic progenitor (*Cd177*+, *Ngp*+) (**Figure 7A, Suppl. Figures 7B-C** and **Suppl. File 7**) To determine whether the p.T73N mutation affects the dynamics of HSPCs differentiation, we used Monocle 3 to perform trajectory analysis on HSPCs populations (after exclusion of two small residual populations of differentiated neutrophils and T cells). The origin of the trajectory was assigned to a more primitive and quiescent HSC (cluster 5). This analysis showed that along the myeloid maturation trajectory, the HSPCs in *MycT73N/+* mice exhibited an aberrant branch decision that delayed the progression of the GMP-like cells (cluster 1) along the differentiation path towards granulocytopoietic progenitor (cluster 10) (**Figure 7B**). Simultaneous examination of *Myc* expression along the same trajectory (**Suppl. Figure 7D**), showed low/absent *Myc* mRNA levels in the stem cell clusters (cluster 5 and 4), upregulation of *Myc* in myeloid progenitors (cluster 1 and 6), followed by downregulation of *Myc* upon progression towards more mature stages (cluster 10), a behavior that is consistent with previous reports (Wilson et al. 2004; Delgado and Leon 2010). *MycT73N/+* mice exhibited a comparable pattern of *Myc* expression along the same trajectory, which exposed how the delay in progression along the maturation trajectory occurred at the stage of initial *Myc* mRNA upregulation, within the GMP-like population (cluster 1), potentially justifying the preferential expansion of GMP+ cells in *MycT73N/+* mice.

**Figure 7.**
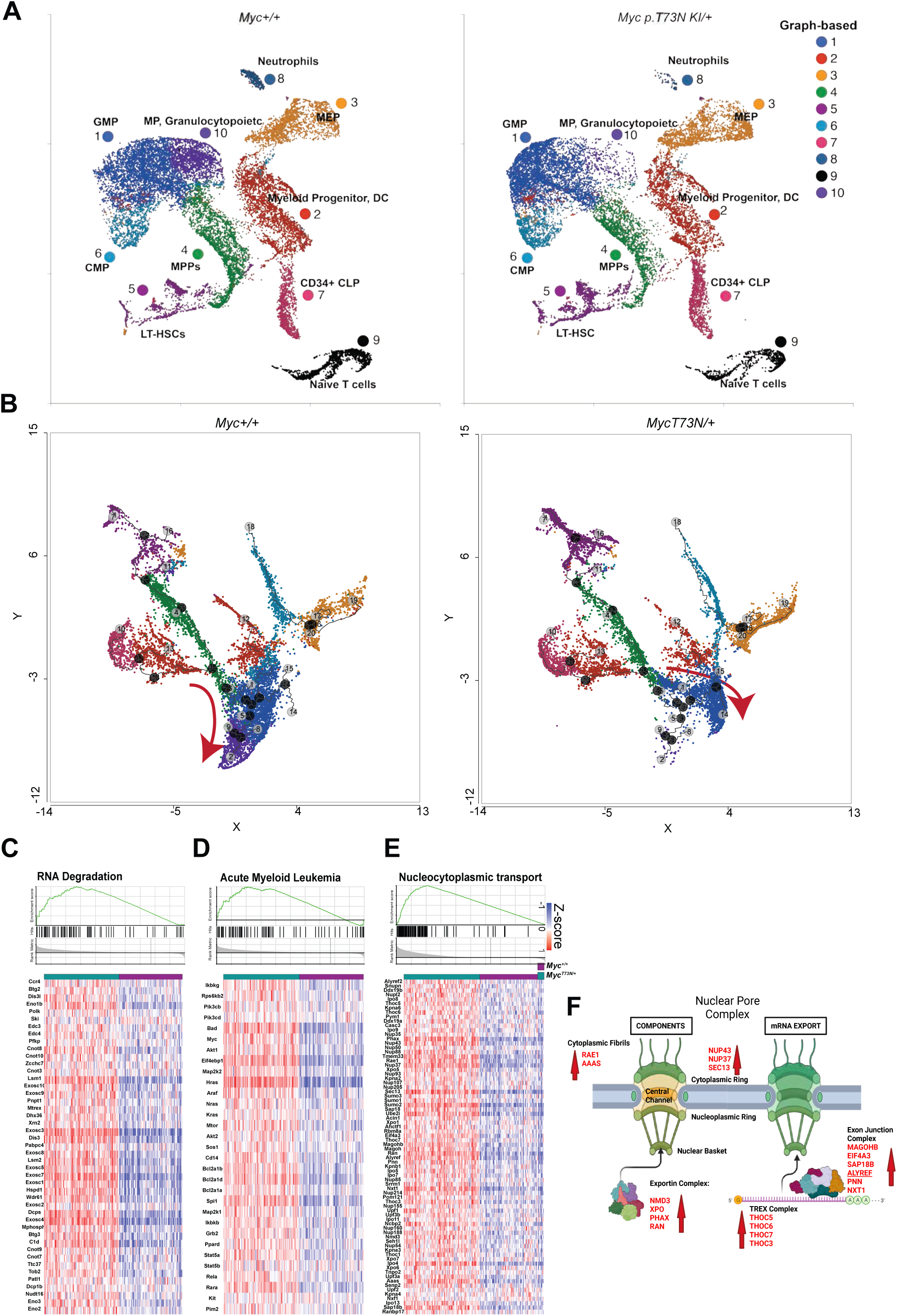
Transcriptomics studies of HSPCs derived from *MycT73N/+* and wildtype littermate mice. (**A**) UMAP projection of 36085 cells derived from 6-months old *MycT73N/+* mice and littermate controls (n=2 per genotype). Colors reflect graph-based clustering, based on transcriptomic similarities amongst the cells, as indicated on the figure (GMP= granulocyte-macrophage; DC=dendritic cell progenitor; MEP= megakaryocyte erythroid progenitor; MPP= multipotent progenitor; LT-HSC= long-term hematopoietic stem cell; CMP= common myeloid progenitor; CLP= common lymphoid progenitor; MP= myeloid progenitor) (**B**) Trajectory analysis of clusters 1-7 and 10 in *Myc^+/+^* (left panel) and *MycT73N/+* (right panel). (**C-E**) Significantly enriched gene sets in GMP- like cells of *MycT73N/+* versus *Myc^+/+^* mice. Gene set enrichment analysis plots are at the top and heatmaps of leading-edge genes on the bottom. Normalized Enrichment Scores is 2.45 for (**C**), 2.21 for (**D**) and 2.56 for (**E**), FDR=0 for all. (**F**) Schematic summary of the nucleocytoplasmic transport components’ transcripts that are upregulated >2 folds in *MycT73N/+* versus *Myc^+/+^*.

Gene set enrichment analysis of (GSEA) of *MycT73N/+* vs *wildtype* GMP-like HSPCs showed enrichment in RNA degradation, acute myeloid leukemia, and nucleocytoplasmic transports gene sets (**Figures 7C-F**; normalized enrichment scores of 2.45, 2.21 and 2.56 respectively, FDR=0). *Nmd3*, *Xpo6* and *Thoc5* were amongst the leading-edge genes upregulated in the nucleocytoplasmic pathway, overlapping with the proteomic results presented in **Figure 4B**. Ribosome Biogenesis, Pyrimidine Metabolism and Signaling pathways regulating stem cells pluripotency were other, highly significant enriched pathways in *MycT73N/+* GMP-like cells (**Suppl. Figures 7E-F**). Conversely, and in keeping with the RNA-seq results from the overexpression models, hematopoietic cell lineage and immune pathways were enriched in GMP-like HSPCs of wildtype mice (**Suppl. Figure 7G**). Overall, these data indicate that the distribution of the HSPCs along the differentiation trajectory is altered between *MycT73N/+* and *wildtype* mice. *MycT73N/+* HSPCs exhibit a maturation block which coincides with the upregulation of *Myc* mRNA expression at the transition between MPP-like and GMP-like myeloid progenitor. The GMP population is expanded in *MycT73N/+* mice and displays abnormal transcriptional patterns in several genes in the mRNA degradation and nuclear export pathways.

## DISCUSSION

The detection of recurrent missense *MYC* mutations in the dominant clone of AML patients has presented us with the unique opportunity to examine the mechanisms of myeloid transformation mediated by MYC. In this study, we employed several orthogonal methods to show that MBI mutant MYC proteins can transform healthy myeloid GMP progenitors. The molecular phenotypes of this transformation partially overlap with those observed in a wildtype MYC overexpression model. However, important differences exist that are unique to the mutant proteins and therefore provide mechanistic clues about how these mutations may contribute to AML development.

Similar to wildtype MYC expression, MBI *MYC* mutations promote transcriptional programs that skew HSPCs balance towards self-renewal at the expense of myeloid differentiation. However, our study uncovered unique and previously unrecognized properties of MBI MYC^MUT^ proteins, notably that these mutants display enhanced leukemogenic potential compared to overexpression of wild type MYC. This increased leukemogenicity appears to be in part related to the mutant proteins’ heightened ability to enhance nuclear export and decay of transcripts involved in cell cycle, apoptosis, and immune regulation, through the modulation of nuclear export complex components. These findings highlight a previously underappreciated role for aberrant nucleocytoplasmic transport in the pathogenesis of AML, which appear to selectively accelerate the decay of specific transcripts. This selectivity may be linked to the cargo specificity of the RNA binding proteins and exportins preferentially induced by MBI MYC^MUT^ expression or could be related to the intrinsic properties of some mRNAs. For instance, changes in mRNA stability have been shown to shape the transcriptome in cancer (Perron et al. 2022) and studies on mRNA turnover dynamics has revealed that genes associated with cellular identity are inherently less stable than RNA processing, RNA binding, and housekeeping transcripts (Kawata et al. 2020; Li et al. 2023). Therefore, an alternative possibility is that the changes in nuclear export may affect transcripts more broadly but manifest transcriptionally as a reduced expression only of those genes with inherently shorter half-lives.

Another striking difference that we found between wild type MYC and MBI missense *MYC* mutations is that although these mutants display increased MYC protein levels and chromatin localization compared to endogenously expressed wildtype MYC, they show decreased occupancy at MYC’s binding sites genome-wide compared to similarly increased, oncogenic expression of wildtype MYC. This became apparent solely because we directly compared the binding profiles of oncogenic levels of MYC wildtype and MBI MYC mutant expression. This finding led to the subsequent observation that, while MBI missense *MYC* mutations did not directly influence the nuclear localization signal sequence of MYC, they result in a distinct and prominent accumulation of MYC within the cytoplasm of hematopoietic stem and progenitor cells. This phenotype was also prominent in the knock-in mouse, where we detected a strong immunofluorescent signal from the MYC protein in the cytoplasm. Furthermore, the cytoplasmic accumulation of mutant MYC preceded the development of the cancer phenotype, as evidenced by its detection in phenotypically healthy *MycT73N/+* mice. Of note, transcriptionally inactive MYC fragments (*Myc-nick*) can form in the cytoplasm through calpain-mediated proteolysis (Conacci-Sorrell et al. 2010) of full-length MYC. These cytoplasmic MYC fragments are observed in normal cells but are prominent in tumors, where they promote tumor cell survival through autophagy induction and apoptosis inhibition (Conacci-Sorrell et al. 2014b). We speculate that the perturbation of nucleocytoplasmic transport components in favor of nuclear export, may offer a partial explanation for the preferential cytoplasmic accumulation of MBI MYC^MUT^. In relation to this, it has been reported that post-translational modifications that increase MYC protein stability shift MYC localization from the nucleoplasm to the nuclear pore basket (Su et al. 2018), suggesting the possibility that stabilized form of MYC proteins may be preferentially exported.

Although the full implications of the cytoplasmic accumulation of these MBI MYC mutants are unclear at this juncture, it aligns with a broader pattern seen for many oncoproteins (Lee et al. 2013; Wang and Li 2014), where mislocalization occurs in tumor cells compared to their normal counterparts. This may lead to the acquisition of functions that could, in turn, contribute to their oncogenic potential. In support of a potential causal relationship between the cytoplasmic MYC localization and the promotion of tumorigenesis, a histological study that explored MYC staining patterns in diffuse large B cell lymphoma (DLBCL) showed that in DLBCLs cases that lack the traditional tumor-initiating *MYC* translocations, the MYC protein is primarily localized to the cytoplasm in 95% of the cancer cells and equally localized to the nucleus and cytoplasm in the remaining 5% of the cases, but it is never localized exclusively in the nucleus (Ruzinova et al. 2010). Other immunohistochemical studies have observed cytoplasmic staining for Myc in various tumors (Bai et al. 1994; Pietilainen et al. 1995). These observations suggest a complex and intriguing interplay between MYC levels, post-translational modification, cellular localization, and the acquisition of non-transcriptional functions that may be critical to cell transformation.

Our study has also revealed that MBI MYC^MUT^ leukemogenic activities can be cell-type specific, and result from the preferential expansion of committed myeloid progenitor cells transcriptionally similar to a GMP cell. While deregulation of MYC has been shown to drive malignant transformation of virtually any cell type, the use of the *MycT73N/+* mouse model underscored that in situations where MYC expression is not enforced by specific promoters, the effects on cell expansion and tumor initiation predominantly impact those cell states that first manifest reliance on MYC expression. It is important to note that although histological examination of the hematopoietic tissues showed the presence of abnormal biphasic-appearing infiltrates in the spleen and lymph nodes of leukemic MycT73^KI/+^ mice, we could not detect any expansion of lymphoid populations in the bone marrow of pre-leukemic mice by scRNA-seq. Similarly, in the tumors we analyzed using flow cytometry, we only detected expansion of myeloid progenitors in the bone marrow. This observation raises the possibility that the histological findings of a biphasic infiltrate in the spleen and lymph nodes point to a source of lymphoid transformation which may originate outside of the bone marrow in other hematopoietic tissues, such as the thymus, spleen, or lymph nodes. Another possibility is that the extramedullary biphasic infiltrate may represents an abnormal population of the same lineage.

AML is a genetically heterogeneous disease, where seemingly unrelated genetic alterations ultimately result in similar cellular phenotypes. This implies that different genetic lesions may converge on common oncogenic pathways. Our study suggests that the dysregulation of RNA partitioning and metabolism, through modulation of nuclear pore complex components by *MYC* mutations, could play an important role in accelerating the transition of hematopoietic progenitors from healthy to a pre-malignant state.

## METHODS

Immunoblotting, Immunohistochemistry, Immunofluorescence, chromatin immunoprecipitation followed by sequencing, RNA sequencing, RNA half-life determination, and proteomic methods are fully detailed in the Supplementary Appendix.

### Plasmid constructs

Retroviral vectors expressing the cDNA of wild-type *c-Myc* and *GFP* in the murine stem cell virus backbone were purchased from Addgene (*MSCV-Myc-IRES-GFP*, Plasmid #18770). Doxycycline-inducible constructs were obtained by cloning *c-Myc* cDNA into *GC385-S* backbone, obtained by modifying the *TtRMPVIR* plasmid (Addgene, Cambridge, MA, USA, plasmid #27995). *Myc^MUT^* (A59V, T73A, T73N, P74L, P74Q, E137Q) were generated at Genewiz (Azenta Life Sciences) through site-directed mutagenesis. Empty backbones vectors (*MSCV-IRES-GFP* and *GC385-S*) were used as controls. All plasmid sequences were verified by Sanger sequencing. Plasmid sequences are available in the Supplementary Appendix.

### Generation of viruses and cells used for *in vitro* and *in vivo* experiments

GP2-293T cells (Takara Bio,631458) were co-transfected with EcoPack packaging plasmid and the *MSCV-IRES-GFP* (Empty backbone or *Myc^WT^* or *Myc^MUT^*) retroviral plasmids using TransIT®-LT1 (Mirus, MIR 2300) according to manufacturer instructions.

Retroviral supernatant was collected two and three days after transfection of GP2-293T cells and concentrated onto wells of a 6 well plates coated with Retronectin (Takara Bio, 5 ug/ml) by spinning at 2500g for 90 minutes at 32° C. Viral supernatant was aspirated from the wells, and Lin^low^ mouse bone marrow cells were added to the wells and transduced by spinning at 280g for 7 minutes at 32° C. Transduction was repeated the following day. Cells were sorted for GFP the day after the second transduction and used in subsequent experiments.

Constitutive *Myc* constructs were used for *in vitro* growth assessment, half-life protein determination and *in vivo* leukemia generation.

The same protocol was used to generate Doxycycline-inducible (Tet-On) viruses. Doxycycline hydrochloride was purchased from Sigma (cat # D3447). Doxycycline-inducible *Myc* constructs were used for *in vitro* growth assessment, *in vivo* leukemia generation and ChIP-sequencing, when comparable expression of both MYC^WT^ and MBI MYC^MUT^ protein were warranted.

Lin^low^ enriched progenitors were obtained by isolating whole bone marrow from wild-type C57BL/6 mice (4 to 6 weeks old) using standard flushing technique. Cells were lineage depleted on an autoMACS using the Direct Lineage Cell Depletion Kit (Miltenyi Biotec, 130-110-470) according to the manufacturer’s protocol. Lin^low^ cells were transduced on day +1 and +3 after lineage depletion. Transduced cells were sorted for GFP on Day +5 post lineage depletion and used for all *in vitro* and *in vivo* experiments. During this process, cells were maintained in transplant media (RPMI, 15% FBS and 1% Penicillin/Streptomycin supplemented with murine Stem cell factor (SCF 100ng/ml), FMS related tyrosine kinase 3 (FLT3 ligand 50ng/ml), Thrombopoietin (TPO 10ng/ml), and Interleukin 3 (IL-3 6ng/ml).

### Flow cytometry and Cell Sorting

Bone marrow, spleen, liver, thymus, lymph nodes and peripheral blood cells were isolated post-mortem using standard techniques. Red blood cell lysis was performed when appropriate using Ammonium Chloride Potassium (ACK) buffer. Cells were stained with flow antibodies against various cell-surface markers as detailed in the Supplementary Appendix. Acquisition of flow samples was performed using the ZE5 Yeti flow analyzer (Bio-Rad) and data was analyzed using FlowJo v.10. Results were visualized using GraphPad Prism v9.1.2. Cell sorting was performed on a modified Sony Synergy SY3200 (Sony Biotechnology, San Jose, CA).

### Mouse Husbandry

C57Bl/6 and B6.SJL mice were used as donors for bone marrow cells and as recipients of transplant experiments and were purchased from The Jackson Laboratory. Mice were fed either normal mouse chow, or chow supplemented with doxycycline (GATEWAY Lab Supply / Urban Feed and Supply Co.) to activate the expression of the dox-inducible *Myc^WT^* or *Myc^MUT^* vectors. The Dox dose in the chow was predetermined to produce equivalent expression of the transgene of interest as previously described (Shirai et al. 2015). All mouse work was done in accordance with Institutional Guidelines and approved by the Animal Studies Committee at Washington University in Saint Louis, under mouse protocol #21-0055.

### *MycT73N/+* mice generation

The *Myc p.T73*N mouse was created on a C57Bl6 background using reagents designed and validated by and genotyped at the Genome Engineering & Stem Center (GESC@MGI) at Washington University in St. Louis. Details regarding mouse generation and genotyping are provided in the Supplementary Appendix.

### Statistical analysis and Analytic Software used for analyses

Statistical differences between control and experimental conditions for *in vitro* and *in vivo* experiments were determined with a two-tailed, non-parametric student t-test, or with one-way ANOVA when comparing more than two conditions. For transplant studies, the statistical comparison for survival was performed using the Kaplan-Meier estimator. All statistical analyses were conducted using Prism. Statistical analyses and graphical outputs for ChIP-sequencing, transcriptomic and proteomic analyses were performed in Partek Flow® (version 10.0.22.1204). Experimental schemas were generated using BioRender.com. Additional software used for single cell RNA-sequencing included Altanalyze (version 2.1.3., http://www.altanalyze.org).

## Acknowledgments

We thank Dan Schweppe and Michael Savio of the Siteman Flow Cytometry Core for their skilled sorting expertise. We thank the Morphology Core at Washington University for embedding, cutting, and staining the mouse-derived tissues. We are grateful to the Genome Technology Access Center at the McDonnell Genome Institute at Washington University School of Medicine for help with genomic analysis. The Center is partially supported by NCI Cancer Center Support Grant #P30 CA91842 to the Siteman Cancer Center. We thank the GESC@MGI and Transgenic, Knockout and Microinjection Core at Washington University in St. Louis for creating the *MycT73N* KI mouse. We are grateful to Petra Erdmann-Gilmore, Dr. Yiling Mi, Alan Davis and Rose Connors for their expert technical assistance with proteomic. The proteomic experiments were performed at the Washington University Proteomics Shared Resource (WU-PSR), R Reid Townsend MD.PhD., Director, Drs. Robert W. Sprung and Qiang Zhang, PhD., Co-Directors. The WU-PSR is supported in part by the WU Institute of Clinical and Translational Sciences (NCATS UL1 TR000448), the Mass Spectrometry Research Resource (NIGMS P41 GM103422) and the Siteman Comprehensive Cancer Center Support Grant (NCI P30 CA091842). This work was supported by a National Cancer Institute (NCI) K08 grant (CA252632), a Specialized Program of Research Excellence in Acute Myeloid Leukemia grant (P50 CA171963) and a Gabrielle’s Angel Foundation for Cancer Research Scholarship (P23-00364) to Dr. Ferraro.

## Authorship Contribution

N.A. and K.C. performed experiments, collected, analyzed, interpreted the data, and drafted portions of the manuscript. M.B.R. performed histological analyses and edited the manuscript. K.M., R.D, C.Y. and A.K. performed experiments and collected data. R.S., P.E.-G., Y.M., and R.R.T. performed the mass spectrometry, the relative data processing and edited the manuscript. A.G. processed, QA/QC, and supervised the bioinformatic data. S.M.S and D.S. participated in the analysis interpretation and critical manuscript revision. FF designed the study, generated reagents, performed and supervised the experiments, collected, analyzed, interpreted the data, and wrote the manuscript.

All authors have read and agreed to the submitted version of the manuscript.

